# Tel1 activation by the MRX complex is sufficient for telomere length regulation but not for the DNA damage response in *S. cerevisiae*

**DOI:** 10.1101/684522

**Authors:** Rebecca Keener, Carla J. Connelly, Carol W. Greider

## Abstract

Previous models suggested that regulation of telomere length in *S. cerevisiae* by Tel1(ATM) and Mec1(ATR) parallel the established pathways regulating the DNA damage response. Here we provide evidence that telomere length regulation differs from the DNA damage response in both the Tel1 and Mec1 pathways. We found that Rad53 mediates a Mec1 telomere length regulation pathway but is dispensable for Tel1 telomere length regulation, whereas in the DNA damage response Rad53 is regulated by both Mec1 and Tel1. Using epistasis analysis with a Tel1 hypermorphic allele, Tel1-hy909, we found that the MRX complex is not required downstream of Tel1 for telomere elongation but is required downstream of Tel1 for the DNA damage response. Since models that invoke a required end processing event for telomerase elongation are primarily based on the yeast pathways, our data call for a re-examination of the requirement for telomere end processing in both yeast and mammalian cells.

## Introduction

Telomere length regulation is critical for cell viability and disruption of length homeostasis leads to disease (STANLEY AND ARMANIOS 2015). Telomerase adds telomere repeats onto chromosome ends and redundant pathways tightly regulate this addition. In humans, decreased telomerase activity causes short telomere syndromes (ARMANIOS AND BLACKBURN 2012) while telomerase activation promotes cancer growth (GREIDER 1999). Thus, understanding the feedback pathways for maintaining telomeres is critical to understanding disease. The Ataxia Telangiectasia-Mutated (ATM) and Ataxia-Telangiectasia and Rad3-related (ATR) checkpoint kinases play a role in both sensing damage and maintaining telomeres around an equilibrium point in yeast (RITCHIE *et al*. 1999) and in mammalian cells (LEE *et al*. 2015; TONG *et al*. 2015; DE LANGE 2018) yet their underlying mechanisms remain unclear.

In *Saccharomyces cerevisiae* multiple independent pathways control telomere length. Tel1, the ATM homolog, and Mec1, the ATR homolog, regulate partially redundant pathways that affect both telomere length and the DNA damage response. *TEL1* and *MEC1* mutations shorten telomeres, although *TEL1* deletion has a greater effect on telomere shortening than *MEC1* mutants. However, the double mutant has an additive effect on telomere shortening, suggesting the kinases regulate parallel pathways (RITCHIE *et al*. 1999). While hypomorphs in *MEC1* show clear telomere shortening, *MEC1* deletions have led to some conflicting conclusions in the literature (DI DOMENICO *et al*. 2014). *MEC1* is an essential gene as *mec1Δ* cells are inviable because they are not able to activate dNTP production. *mec1Δ* cells survive only with co-deletion of either *SML1* or *CRT1*, and in *mec1Δ sml1Δ* or *mec1Δ crt1Δ* cells, dNTP production is increased. *mec1Δ sml1Δ* and *mec1Δ crt1Δ* have different effects on telomere length which has been attributed to the fact that *SML1* and *CRT1* regulate different pathways of dNTP production (MAICHER *et al*. 2017). Mec1 and Tel1 also both play a role in the DNA damage response, where Mec1 has the greater effect. *TEL1* deletion on its own does not show DNA damage sensitivity, but strains with mutations in both *TEL1* and *MEC1* show higher sensitivity to DNA damage than *MEC1* mutation alone (MORROW *et al*. 1995). These experiments indicate that *TEL1* and *MEC1* have roles in both the DNA damage response and telomere length regulation.

The distinct effects of *tel1Δ* and *mec1Δ* on telomere length and the DNA damage response suggest Tel1 and Mec1 may have different critical substrates. Both Tel1 and Mec1 phosphorylate S/T-Q motifs (KIM *et al*. 1999). This identical phosphorylation motif has made identifying unique substrates of each kinase challenging. Mass spectrometry approaches have identified specific Tel1 and Mec1 substrates, in addition to shared substrates (BASTOS DE OLIVEIRA *et al*. 2015), but the biological consequences of these phosphorylation events remain unclear.

While hundreds of Tel1/Mec1 substrates have been identified, those that are critical for the DNA damage response and telomere length have not been defined. We investigated the role of two known substrates: Rad53 and the MRX complex. Both candidates were reported to be phosphorylated by Tel1 and/or Mec1 in response to DNA damage (D’AMOURS AND JACKSON 2001; NAKADA *et al*. 2003b; SMOLKA *et al*. 2007; ALBUQUERQUE *et al*. 2008; BASTOS DE OLIVEIRA *et al*. 2015; LAVIN *et al*. 2015). In the DNA damage response, both Tel1 and Mec1 phosphorylate Rad53, activating it’s kinase activity (Figure 1A)(NAKADA *et al*. 2003b). However, Tel1 phosphorylation is considered less important relative to Mec1 phosphorylation of Rad53 (USUI *et al*. 2001). In response to a double-strand break, Tel1 interaction with the MRX complex activates Tel1 kinase activity and Tel1 subsequently phosphorylates the MRX complex, in addition to other substrates (LEE *et al*. 2013; BASTOS DE OLIVEIRA *et al*. 2015). The MRX complex resects the double-strand break to facilitate repair and several studies indicate that Tel1 modulates this process (LAVIN *et al*. 2015). In this model the MRX complex can be considered both upstream and downstream of Tel1 for the DNA damage response (Figure 1A). Several studies have suggested that similar regulatory events occur at the telomere (TSUKAMOTO *et al*. 2001; LARRIVEE *et al*. 2004; VISCARDI *et al*. 2007; BONETTI *et al*. 2009), although specific mechanisms are not well established.

**Figure 1.**
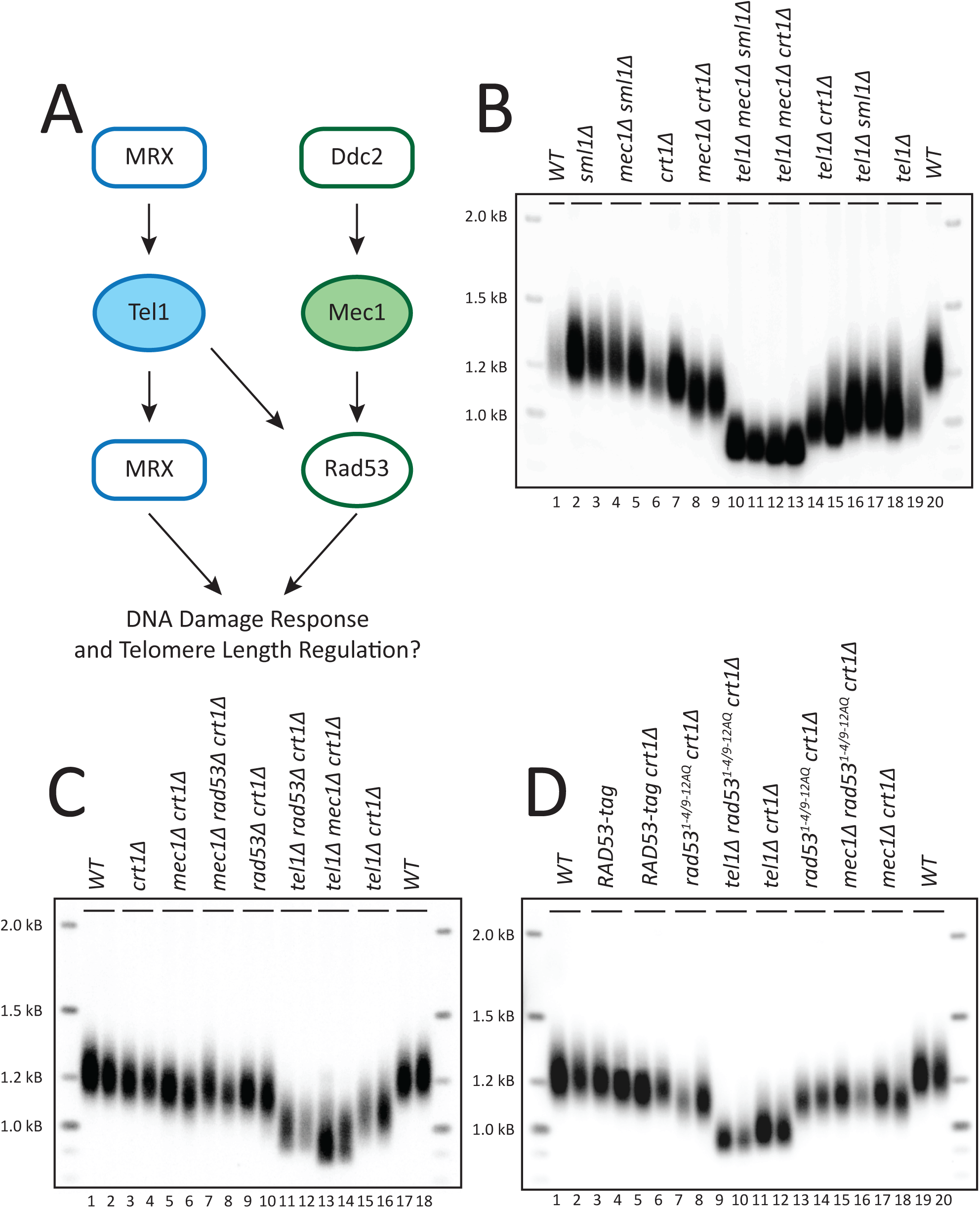
Rad53 is in the Mec1 telomere length regulation pathway. (A) Diagram representing a simplified, current understanding of Tel1/Mec1 pathways in the DNA damage response. (B)-(D) Southern blot analysis of telomeres from segregants with the indicated genotype. Two independent, haploid segregants were assayed for each genotype. (B) Haploid cells were passaged on solid media for approximately 120 population doublings to decrease telomere length heterogeneity. Segregants are from JHUy937-1. (C) Segregants are from yRK6002 and yRK6003. (D) Both Rad53 and Rad53^1-4/9-12AQ^ are epitope tagged with a 3xFLAG tag. Haploids were passaged for approximately 100 population doublings. Segregants are yRK6008-1, yRK6008-2, yRK6009-1, yRK6009-2, yRK6010-1, yRK6010-2, yRK6011-1, yRK6011-2, yRK6012-1, yRK6012-2, yRK6013-1, yRK6013-2, yRK6014-1, yRK6014-2, yRK6015-1, and yRK6015-2.

In this study we used mutagenesis and epistasis analysis to show that *RAD53* is in the *MEC1* telomere length pathway. In addition, epistasis analysis showed the MRX complex acts both upstream and downstream of Tel1 in the DNA damage response, as characterized by others. However, strikingly, the MRX complex is only required upstream of Tel1 in telomere length regulation. Therefore, while the MRX complex is required to activate Tel1 kinase activity, it is not required for telomere resection. These findings demonstrate that the regulation of these proteins in the DNA damage response are distinct from their regulation in telomere length maintenance and challenge the assumption that a telomere must be resected by the MRX complex for telomere elongation by telomerase.

## Materials and Methods

### Molecular Cloning

Each plasmid was constructed using Gibson Assembly (GIBSON 2011), for detailed explanations of cloning strategies see File S1. Primers were designed using Snapgene software (GSL Biotech), products were amplified with Phusion HS II DNA polymerase (Thermo Fisher F549), and Gibson Assembly Master Mix (New England Biolabs E5510) was used according to the New England Biolabs (NEB) recommended protocol. All restriction enzymes and NEB5*α* competent cells (NEB C2987H) are from NEB. Plasmids were prepared using QIAprep Miniprep Kit (Qiagen 27106) and all sequencing was performed using the Sanger method.

### Site-directed mutagenesis

S/T-Q mutations and *TEL1-hy909* mutations were introduced by site-directed mutagenesis using primers designed by PrimerX.org. Primer sequences are listed in Table S3. In each case, the plasmid was amplified using PfuTurbo (Agilent 600252). The product was *Dpn*I-treated, ethanol precipitated, and transformed into DH5*α* cells (Thermo Fisher 18265017). Clones were isolated and sequence verified.

### Yeast culturing and transformation

Yeast culturing, transformation, and sporulation were conducted as described in (GREEN AND SAMBROOK 2012). Briefly, transformation was carried out on 50 mL of logarithmically cultured cells treated with 0.1 M Lithium Acetate (LiAc, Sigma L6883). DNA was added to a 50 µL aliquot of cells in addition to 50 µg boiled fish sperm carrier DNA (Roche 11467140001). Cells were equilibrated at 30° for 10 minutes after which 0.5 mL 40% Polyethylene glycol (PEG_4000_, Sigma P4338) containing 0.1 M LiAc was added. Cells were incubated at 30° for 30 minutes then heat shocked at 42° for 15 minutes. Transformed cells were washed with sterile water and plated on the appropriate selective media. In cases where an antibiotic selectable marker was used, cells were recovered in 1 mL Yeast-extract Peptone Dextrose (YPD) at 30° for 3-4 hours before plating. One-step integration was used for all integrated constructs. After transformation, integration at the desired locus was confirmed by junction Polymerase Chain Reaction (PCR). In cases where mutations were introduced, the region was amplified using Phusion HS II DNA polymerase the amplicon was purified using AMPure beads (Beckman Coulter A63881), and sequenced to confirm the presence of mutations *in vivo*.

### Passaging yeast

Because we initiate our experiments with strains that are heterozygous for multiple alleles of interest, dissection of twenty tetrads often yields all combinations of the alleles. This makes it possible to obtain experimental and control samples in parallel. Treating all haploids in parallel is critical for evaluating telomere length as telomere length can be sensitive to differences in culturing time. In cases where the diploid genotype did not allow isolation of an important control, a second diploid, from which that control can be segregated, was dissected in parallel. After tetrad dissection and replica plating to identify segregants of interest, haploids were streaked to single cell on a YPD plate. This streak was designated as the first passage, after which these cells are estimated to have undergone 40 population doublings. Subsequently, cells were re-streaked repeatedly to increase the number of cell divisions. Each passaged plate was incubated for 48 hours at 30° after which cells were picked from the streak dilution and re-streaked to single cell again on a fresh plate. Each passage is estimated to require approximately 20 population doublings. Therefore, a strain passaged five times undergoes a total of approximately 120 population doublings. At the desired number of passages, cells were inoculated into a 5 mL liquid YPD culture and grown at 30° overnight or until saturated. Cell pellets were saved at −20° until all time points had been collected, at which time genomic DNA was extracted. Images of the plates at specific time points were taken to document potential growth defects.

### *MRX-tag* and *mrx-18A* strain construction

A yRK1006 *MRE11-3HA-URA3* segregant was mated to a yRK2040 *RAD50-G6-V5-LEU2* segregant, yielding yRK1. The *XRS2-13myc-hphMX4* construct (pRK1028) was transformed into yRK1, yielding yRK60. A yRK60 segregant with all three *MRX-tag* components was mated to a *mec1Δ sml1Δ* haploid (JHUy816 segregant) to yield yRK79 and yRK80. The *mrx-18A* parental diploids were constructed in a manner parallel to the *MRX-tag* parental diploids. A yRK1052 *mre11-4A-3HA-URA3* segregant was mated to a yRK2082 *rad50-10A-G6-V5-TRP1* segregant, yielding yRK26. The *xrs2-4A-13myc-hphMX4* construct (pRK1040) was transformed into yRK26, yielding yRK56. A yRK56 segregant with all three *mrx-18A* components was mated to a *mec1Δ sml1Δ* haploid (yYM242 segregant) to yield yRK81 and yRK83. Mutations were confirmed again in yRK81 and yRK83 by sequencing.

### *rad50S* knock-in using CRISPR/Cas9

gRNA sequences were chosen using the algorithms published by Doench et al. via the Benchling (Biology Software, 2019) interface (DOENCH *et al*. 2016). Two guides, RW 670 and RW 671, were individually cloned into pCAS026 using the strategy described by Anand et al. (ANAND *et al*. 2017). gRNA sequences are listed in Table S3. Double-stranded homology repair templates were amplified (primers RW 674, RW 675) using a 90-mer oligonucleotide as the template. The repair template was designed to both introduce the Rad50S mutation (Lys81Ile) (ALANI *et al*. 1990) and introduce a silent mutation in the Protospacer Adjacent Motif (PAM) sequence. Primer RW 672 was used as the template for the RW 670 guide and primer RW 673 was used as the template for the RW 671 guide. SPRI beads (Beckman Coulter B23318) were used to purify the repair template before transformation. yRK114 haploid cells were cultured for transformation as described above except that no carrier DNA was added and 100 µL of cells were used in the transformation reaction. Cells were transformed with 1 µg of pCAS026 containing the appropriate guide and 4-5 µg of double-stranded repair template. Cells were grown on minimal media without uracil to select for cells that contained the pCAS026 plasmid. The *rad50S* allele was validated by sequencing. yRK2112 was edited using the RW 670 guide while yRK2116 was edited using the RW 671 guide. yRK2113 was transformed with pCAS026 containing the RW 670 guide but was not edited and was used in parallel to serve as a control. Haploids were streaked on minimal media containing 5-Fluoroorotic Acid (5-FOA) (Toronto Research Chemicals F595000) to select against the pCAS026 plasmid followed by five passages on YPD plates before telomere elongation was observed by Southern blot (data not shown).

### Southern blot analysis

Genomic DNA was extracted and used for Southern blot analysis as described previously (KAIZER *et al*. 2015). Briefly, a cell pellet of approximately 50 µL was resuspended in 1x Lysis Buffer (10 mM Tris, pH 8.0, 0.5 M EDTA, pH 8.0, 100 mM NaCl, 1% SDS, 2% Triton X-100) and cells were lysed in the presence of 0.3mm glass beads (Biospec products, Inc. 11079105). Phenol-chloroform (50:50) was added and cells were vortexed for 8 min (Eppendorf mixer 5432). The DNA was ethanol precipitated and resuspended in 40-50 µL TE (10 mM Tris, pH 8.0, 1 mM EDTA) with RNaseA (10 µg/mL) at 37° for one hour or 4° overnight.

Samples were cut with the restriction enzyme *Xho*I and electrophoresed at 47 V (12 mA) on a 1.0% agarose in 1x TTE buffer (20x = 1.78 M Tris base, 0.57 M Taurine, 0.01 M EDTA) for approximately 24 hours. 200 ng of 2-log DNA ladder (NEB N3200) was included for reference. The genomic DNA was transferred to a Hybond N+ membrane (GE Healthcare RPN303B) by vacuum blotting (Boekel Appligene vacuum blotter) for 1 hour at 50 mbar with the gel covered in 10x SSC buffer (1.5 M NaCl, 0.17 M sodium citrate). Once transferred, the DNA was UV crosslinked (Stratagene UV Stratalinker 2400). The membrane was prehybridized in Church buffer (0.5 M Tris, pH 7.2, 7% SDS, 1% bovine serum albumin, 1 mM EDTA) at 65° then *α*^32^P-dCTP-radiolabeled (Perkin Elmer) fragments of the Y’ element and the 2-log DNA ladder were added at 10^6^ cpm/mL and 10^4^ cpm/mL, respectively. The membrane was incubated with the radiolabeled probe overnight, washed in 1x SSC, 0.1% SDS buffer at 65°, imaged with a Storage Phospho Screen (GE Healthcare) overnight and then scanned on a Storm 825 imager (GE Healthcare). The images were copied from ImageQuant (GE Life Sciences) to Adobe PhotoShop CS6 and saved as .tif files. The images were cropped in Adobe PhotoShop CS6 to show only the telomere restriction fragment.

### Western blot

Protein extracts were prepared by Trichloroacetic acid (TCA) extraction (LINK AND LABAER 2011). Samples were resolved on a NuPAGE 3-8% Tris-Acetate gradient polyacrylamide gel (Invitrogen EA0375) in 1x Tris-Acetate running buffer (Invitrogen LA0041) using the Invitrogen NuPAGE system with protein ladder standards (Bio-Rad 161-0374). The gel was transferred by electroblotting to a PVDF membrane (Thermo Fisher IPFL00010) using NuPAGE transfer buffer (20x: 40.8 g Bicine, 52.4 g Bis-Tris, 3.0 g EDTA) at 30 V for 1.5 hours. The membrane was blocked with Odyssey buffer (Li-Cor 927-40000) for 1 hour at room temperature or overnight at 4°. Primary antibodies were diluted in blocking buffer and incubated at room temperature for 1 hour (Sigma Aldrich M2 Flag at 1:1,000; Invitrogen 22C5D8 Pgk1 at 1:6,000; Roche 12CA5 HA at 1:2,000; Invitrogen R960-25 V5 at 1:2,000; Santa Cruz 9E10 c-myc at 1:10,000). The membrane was washed in 1x Tris Buffered Saline with Tween-20 (TBST) buffer (10x TBST: 0.2 M Tris Base, 1.5 M NaCl, 1% Tween-20) before incubation at room temperature for 30 minutes with a Horse Radish Peroxidase (HRP) conjugated secondary antibody (Bio-Rad 1706516 at 1:10,000) in 5% powdered milk (Bio-Rad 170-6404) resuspended in 1x TBST. The membrane was washed in 1x TBST and then incubated with Forte HRP substrate (Millipore WBLUF0100) followed by imaging on ImageQuant LAS 4000 mini biomolecular imager (GE Healthcare). The images were copied from ImageQuant (GE Life Sciences) to Adobe PhotoShop CS6 and saved as .tif files.

### Mutagen challenge

Strains of interest were inoculated to an initial OD600 of 0.15-0.25 in 8-10 mL YPD. Once the density reached an OD600 of 0.5-0.6, the culture was split into untreated and treated samples. 4-nitroquinoline (Sigma N8141) was resuspended in acetone at a stock concentration of 1 mM. 15U Bleomycin (Fresenius Kabi C103610) was dissolved in 10 mL sterile water and used as a 1 mg/mL stock. Hydroxyurea (US Biologicals H9120) was resuspended in sterile water at a stock concentration of 1 M. Methyl methanesulfonate (MMS) (Sigma 129925) was treated as 100%. The appropriate chemical was added to each treated sample and cultured at 30° with slight agitation for 1-2 hours, as indicated in the figure legends. Untreated samples were cultured in parallel. Cell pellets of equal density were collected for each sample based on the OD600. The size of the cell pellet varied between experiments from 0.6 OD to 8.0 OD, as indicated in the figure legends. Each pellet was resuspended in 1 mL YPD and serially diluted 1:5 in YPD in a 96-well dish. 4 µL of each dilution was spotted onto a YPD plate and the plates were cultured at 30° for 48 hours before being imaged on a Bio-Rad Gel Doc XR+ Imaging System under white light using Image Lab 6.0.1 software.

### Quantitative MMS survival assay

Freshly grown strains of interest were inoculated to an initial OD600 of 0.2-0.3 in 6 mL YPD. Once density reached 0.5-0.6 OD600, an untreated sample was plated for each strain before cells were treated with 0.01% MMS. 30-minute time points were taken up to 120 minutes for each strain. A Millipore Scepter with 40 µm tips was used to measure cells/mL at each time point. Approximately 500 cells were plated across five YPD plates, with approximately 100 cells per plate, for each strain and at each timepoint. Samples were blinded before plating and the plates were incubated at 30° for 48 hours. Colony forming units were counted for each blinded sample and once all plates were counted, the results were unblinded. At each timepoint the number of colonies was calculated as a proportion: the total number of colonies for that strain at that timepoint relative to the total number of colonies for that strain at the untreated time point (t=0). Data were graphed using Prism 5.0b and the standard error of the mean is shown.

### Plasmid end-joining assay

Cells were cultured and treated as described above for yeast transformation. Once density reached an OD600 of 0.6-0.8 the cells were transformed with 100 ng of *Stu*I-linearized pRS317 (SIKORSKI AND BOEKE 1991), which generates blunt ends, or with 100 ng of circular pRS317. 50 µg boiled fish sperm carrier DNA (Roche 11467140001) was added to both linear and circular transformation reactions. Three replicate transformations were performed for both linear and circular plasmids and for each strain. Cells were plated on minimal media without lysine. After 48 hours of incubation at 30°, colony forming units were counted. The average number of colonies of the three replicates containing circular DNA was calculated for each strain. Each linear DNA transformation was treated as a technical replicate. The number of colonies from each linear DNA transformation plate was normalized to the average number of colonies of circular DNA for that strain. Data were plotted and analyzed in Prism 5.0b. An unpaired two-tailed student *t*-test was performed between samples.

### Data Availability Statement

All strains and plasmids are available upon request. The Reagent Table provides a reference for all genes, strains, software, and many reagents used in this study. Table S1 contains all strains used in this study. Table S2 contains all plasmids used in this study and a brief description of their purpose. Table S3 contains all primers used in this study and a brief description of their purpose. File S1 has a detailed explanation of how all plasmids used in this study were constructed. All supplemental files, including the Reagent Table, have been uploaded to fig**share**.

## Results

### Rad53 regulates telomere length through the Mec1 pathway

Rad53 kinase is a candidate substrate that could mediate telomere length since both Tel1 and Mec1 phosphorylate Rad53 and kinase-dead Rad53 has short telomeres (LONGHESE *et al*. 2000; NAKADA *et al*. 2003b). We used epistasis and mutational analyses to examine whether Rad53 functions in the Tel1 or Mec1 telomere length pathway (Figure 1A). We deleted either *SML1* or *CRT1,* regulators of dNTP pools that were previously shown to suppress the lethality of *mec1Δ* and *rad53Δ (HUANG et al. 1998; ZHAO et al. 1998)*. While it is most common in the literature to use *sml1Δ* to rescue *mec1Δ* lethality, there is evidence that *SML1* deletion can mask telomere length phenotypes (LONGHESE *et al*. 2000). Therefore, we initially compared the effects of deleting *sml1Δ* or *crt1Δ* on telomere length in *tel1Δ* or *mec1Δ* mutants.

All of our experiments were carried out in haploid yeast, however, to avoid telomere length changes that can occur with long term propagation of haploids, we standardly generated fresh haploids by sporulating heterozygous diploids (see methods). We generated diploid strains that were heterozygous for *TEL1/tel1Δ MEC1/mec1Δ SML1/sml1Δ* and *CRT1/crt1Δ* and then sporulated to obtain haploids with specific mutant combinations. While *sml1Δ* and *mec1Δ sml1Δ* cells had telomeres similar to wild-type cells, *crt1Δ* cells had slightly shorter telomeres (Figure 1B, compare lanes 2-5 to lanes 6-7) and *mec1Δ crt1Δ* cells had shorter telomeres than *crt1Δ* cells (Figure 1B, compare lanes 8-9 to lanes 6-7). *mec1Δ crt1Δ* telomeres were similar to the short telomeres reported in *mec1-1* and *mec1-21* cells (data not shown)(RITCHIE *et al*. 1999). *tel1Δ sml1Δ* cells showed short telomeres very similar to *tel1Δ* mutants. However, *tel1Δ crt1Δ* telomere lengths were slightly shorter than *tel1Δ sml1Δ* (Figure 1B, compare lanes 14-15 to lanes 16-17). We conclude that *sml1Δ* masks the short telomere phenotype of *mec1Δ* cells while *crt1Δ* does not, although *crt1Δ* has a mild telomere shortening effect on its own.

To examine whether Rad53 plays a role in the Tel1 or Mec1 telomere length pathway, we generated diploids heterozygous for *TEL1/tel1Δ MEC1/mec1Δ RAD53/rad53Δ* and *CRT1/crt1Δ* and sporulated to obtain specific mutant combinations. We observed that *crt1Δ* suppresses *rad53Δ* lethality, consistent with previous reports (Figure S1B) (HUANG *et al*. 1998). We compared *mec1Δ crt1Δ* telomeres to *mec1Δ rad53Δ crt1Δ* and observed no additive shortening (Figure 1C, compare lanes 7-8 to lanes 5-6), which is consistent with Rad53 functioning in the Mec1 telomere length regulation pathway. In contrast, we found there was additive shortening in *tel1Δ rad53Δ crt1Δ* telomeres compared to *tel1Δ crt1Δ* (Figure 1C, compare lanes 11-12 to lanes 15-16), supporting the conclusion that Tel1 and Rad53 are in different length regulation pathways. However, *tel1Δ rad53Δ crt1Δ* telomeres were slightly longer than *tel1Δ mec1Δ crt1Δ* telomeres (Figure 1C, compare lanes 11-12 to lanes 13-14), suggesting that either Rad53 has functions in both the Tel1 and Mec1 telomere length pathways or that Rad53 acts exclusively in the Mec1 telomere length pathway which also relies on additional Mec1 substrates for telomere length regulation.

### Phosphorylation of Rad53 on S/T-Q motifs regulates telomere length

To examine the role of Rad53 as a Tel1/Mec1 substrate that mediates telomere length, we examined a Rad53 mutant where Tel1/Mec1 S/T-Q phosphorylation motifs were mutated to A-Q. Previous work demonstrated that a subset of the Rad53 S/T-Q motif clusters are critical for Rad53 function in the DNA damage response and that this mutant, *rad53^1-4/9-12AQ^*, could not respond to Mec1 or Tel1 regulation (LEE *et al*. 2003). Both Rad53 and *rad53^1-4/9-12AQ^* were viable when expressed off of a plasmid using the endogenous promoter in *rad53Δ* cells and did not require co-deletion of *SML1* or *CRT1* (Figure S1A and data not shown). We integrated 3xFLAG-tagged *rad53^1-4/9-12AQ^* or 3xFLAG-tagged *RAD53* at the endogenous locus in *TEL1/tel1Δ MEC1/mec1Δ CRT1/crt1Δ* diploid cells to better control the segregation of these alleles. Rad53 and Rad53^1-4/9-12AQ^ were stably expressed, as shown previously (LEE *et al*. 2003)(data not shown). Unlike the plasmid expression system, we noted that integrated *rad53^1-4/9-12AQ^* is lethal unless it co-segregates with *sml1Δ* or *crt1Δ* (Figure S1B). *rad53^1-4/9-12AQ^ crt1Δ* cells had shorter telomeres than *RAD53-tag crt1Δ* cells (Figure 1D, compare lanes 7-8 to lanes 5-6), demonstrating that phosphorylation of Rad53 contributes to telomere length regulation.

We did not observe additive shortening in *mec1Δ rad53^1-4/9-12AQ^ crt1Δ* cells compared to *mec1Δ crt1Δ* or *rad53^1-4/9-12AQ^ crt1Δ* cells (Figure 1D, compare lanes 15-16 to lanes 17-18 and lanes 13-14), consistent with Mec1 targeting these phosphorylation sites. In contrast, we observed additive shortening in *tel1Δ rad53^1-4/9-12AQ^ crt1Δ* cells compared to *rad53^1-4/9-12AQ^ crt1Δ* cells (Figure 1D, compare lanes 9-10 to lanes 7-8), further supporting the conclusion that Tel1 and Rad53 are in different pathways. We observed only a slight shortening in *tel1Δ rad53^1-4/9-12AQ^ crt1Δ* cells compared to *tel1Δ crt1Δ* cells (Figure 1D, compare lanes 9-10 to lanes 11-12). Similar to our earlier findings, this is consistent with either Tel1 and Mec1 phosphorylate Rad53 for telomere length regulation or that there are other Mec1 substrates that are key for telomere length in addition to Rad53.

### Tel1-hy909 requires Rad53 for DNA damage response but not for telomere length regulation

To directly examine whether Tel1 telomere length regulation is Rad53-dependent, we performed epistasis analysis with *rad53Δ crt1Δ* and a Tel1 hypermorphic allele, *TEL1-hy909,* which has increased Tel1 kinase activity, increased DNA damage response function, and long telomeres (BALDO *et al*. 2008). We generated diploids heterozygous for *TEL1/TEL1-hy909 RAD53/rad53Δ* and *CRT1/crt1Δ* and diploids heterozygous for *TEL1/TEL1-hy909 MEC1/mec1Δ* and *CRT1/crt1Δ*. We first examined the DNA damage response in haploid cells by challenging them with a DNA-damaging agent, methyl methanesulfonate (MMS). *mec1Δ crt1Δ* cells were sensitive to DNA damage while *TEL1-hy909 mec1Δ crt1Δ* cells were not (Figure 2A), consistent with previous reports (BALDO *et al*. 2008). This indicates that Tel1 hyperactivity can rescue the DNA damage response in a *mec1Δ* mutant. Like *mec1Δ crt1Δ,* the *rad53Δ crt1Δ* cells were also sensitive to DNA damage (Figure 2B). However, unlike *TEL1-hy909 mec1Δ crt1Δ* cells, *TEL1-hy909 rad53Δ crt1Δ* cells were still sensitive to MMS challenge (Figure 2B). Further, while *TEL1-hy909* was able to rescue *mec1Δ* lethality (Figure S2A), as previously shown (BALDO *et al*. 2008), *TEL1-hy909* was not able to rescue *rad53Δ* lethality (Figure S2B). These data indicate that Rad53 is essential to mediate the Tel1-hy909 DNA damage response.

**Figure 2.**
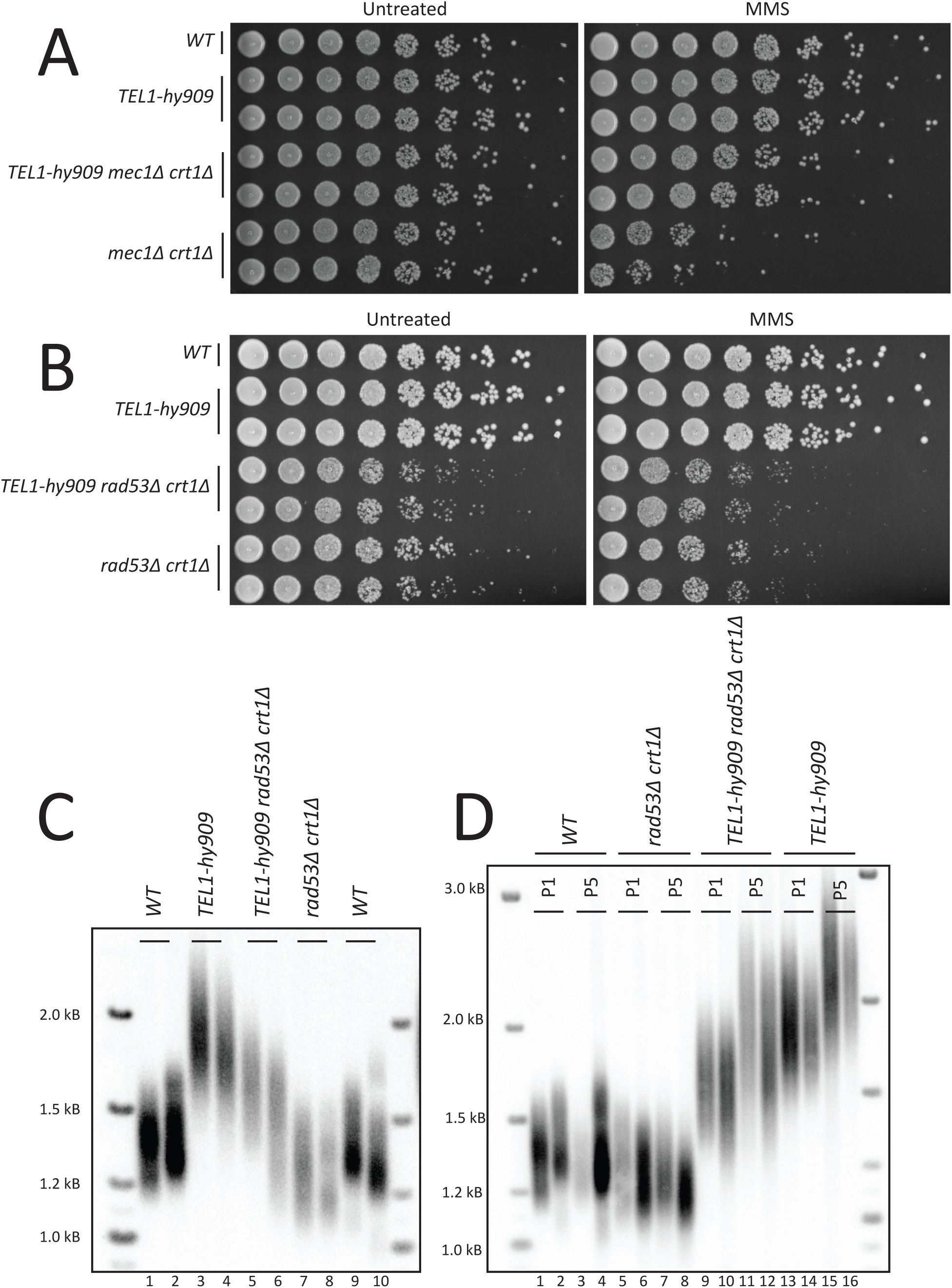
Tel1-hy909 requires Rad53 for the DNA damage response but not telomere elongation. (A)-(B) Yeast dilution series of untreated cells or cells cultured in 0.02% MMS for one hour. The genotype is indicated to the left of the panels. (A) Segregants are from yRK5126 and yRK5127. (B) Segregants are from yRK5028 and yRK5059. (C) Southern blot analysis of telomeres from segregants with the indicated genotype. Two independent, haploid segregants were assayed for each genotype. Because the *TEL1-hy909* hypermorph elongates telomeres in the parental diploid (not shown), we observed increased telomere length heterogeneity across all genotypes in the haploid segregants and observe the wild-type segregant telomeres were longer compared to other Southern blots. Segregants are from yRK5028 and yRK5059. (D) Southern blot analysis of telomeres from segregants with the indicated genotype. Segregants were passaged on solid media for approximately 120 population doublings. Passage number is indicated below the genotype as P1 for the first passage or P5 for the fifth passage. Note that the WT telomeres were longer in the parental diploid due to Tel1-hy909. Segregants are from yRK5028.

We next examined telomere length in *TEL1-hy909* and *TEL1-hy909 rad53Δ crt1Δ* mutants. *TEL1-hy909 rad53Δ crt1Δ* cells had an intermediate telomere length between *TEL1-hy909* and *rad53Δ crt1Δ* (Figure 2C). This indicates that either Rad53 has a partially redundant role in the Tel1 telomere length regulation pathway or that Rad53 and Tel1 function in independent pathways. We reasoned that if Rad53 was important for Tel1-hy909 telomere elongation then telomeres would not further elongate with increased divisions in the absence of Rad53. We passaged *TEL1-hy909 rad53Δ crt1Δ* cells and observed that telomeres elongated with further passages (Figure 2D). This suggests that Rad53 is not required for Tel1 telomere length regulation which is in stark contrast to the Rad53-dependence of Tel1-hy909 for the DNA damage response (Figure 2A). Together, these data suggest that Rad53, likely with other unidentified proteins, functions in the Mec1 telomere length pathway, but is not required in the Tel1 telomere length pathway.

### S/T-Q sites in MRX affect the DNA damage response but not telomere length regulation

Having found that Rad53 is not essential to mediate the Tel1 telomere length response, we next tested whether the MRX complex might be a critical Tel1 substrate. In both yeast and mammalian cells, all three components of the MRX(N) complex are phosphorylated by Tel1/Mec1 (ATM/ATR) (D’AMOURS AND JACKSON 2001; NAKADA *et al*. 2003b; SMOLKA *et al*. 2007; ALBUQUERQUE *et al*. 2008; BASTOS DE OLIVEIRA *et al*. 2015; LAVIN *et al*. 2015). Tel1 and Mec1 have a well characterized and conserved S/T-Q phosphorylation motif (KIM *et al*. 1999). Previous work showed that mutation of all S/T-Q motifs in Xrs2 to A-Q had no effect on the DNA damage response or on telomere length (MALLORY *et al*. 2003). To further probe whether MRX phosphorylation affects function, we mutated all S/T-Q motifs across the entire MRX complex and examined the effect on DNA damage response and telomere length.

Each MRX gene was epitope-tagged, the S/T-Qs were mutated to A-Qs, and the construct was integrated at the endogenous locus (Figure 3A). We generated three strains with each gene individually mutated at all S/T-Q sites, termed *mre11-4A*, *rad50-10A*, and *xrs2-4A*, as well as a strain containing all three mutants, termed *mrx-18A* (see methods). As a control, we generated a strain with all three proteins epitope-tagged but with wild-type coding sequence, referred to as *MRX-tag*. Individually, and in combination with one another, the altered proteins were stable as determined by western blot, indicating that mutations do not affect protein stability (Figure 3B).

**Figure 3.**
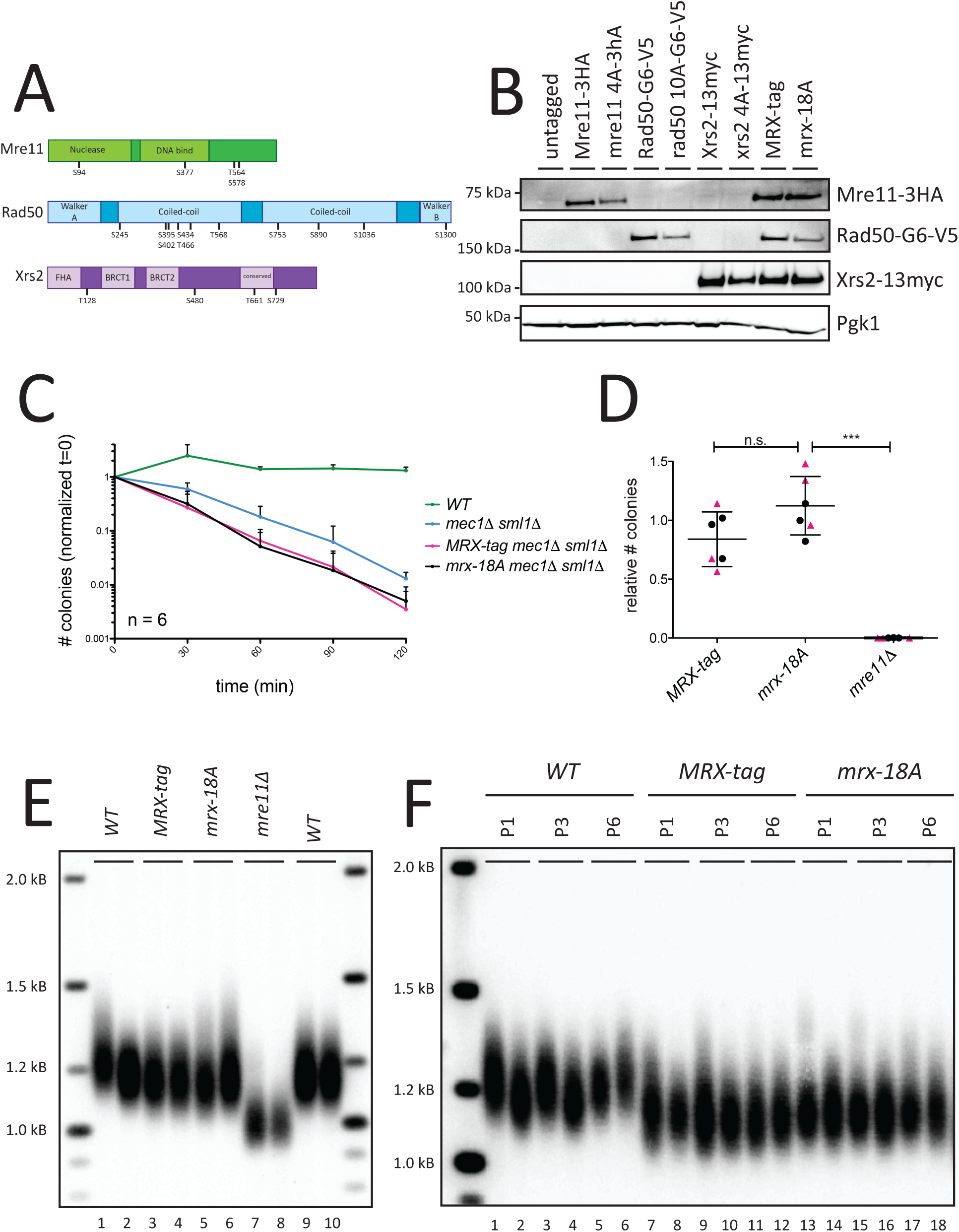
The mrx-18A S/T-Q mutant does not affect the DNA damage response, NHEJ, or telomere length. (A) Domain structure of the MRX complex indicating location of S/T-Q motifs. Mre11 has an N-terminal nuclease domain and a DNA binding domain. Rad50 has an N-terminal Walker A domain and a C-terminal Walker B domain in addition to central coiled-coil domains (LEE *et al*. 2013). Xrs2 has a Forkhead association (FHA) domain followed by two BRCA1 domains and there is a conserved region near the C-terminus (SHIMA *et al*. 2005; BECKER *et al*. 2006). The domain sizes are relatively to scale for each protein and are consistent with NCBI annotation (Mre11: BAA02017.1, Rad50: CAA65494, Xrs2: AAA35220.1). The location of each S/T-Q motif is indicated with the involved S or T residue number indicated. (B) Western blots examining stability of tagged MRX protein components individually or in combination. Samples were run and transferred on duplicate gels simultaneously. The membranes were cut such that each protein could be independently probed. Strains used in western blot are yRK114, yRK106, yRK94, yRK138, yRK139, yRK133, yRK135, yRK102, and yRK90. (C) Proportion of colonies on cells treated with 0.01% MMS over 120 minutes (see methods). Proportion is calculated as the number of colonies at that time point for that genotype relative to the number of colonies for that genotype at t=0. The average and standard error of the mean of six technical replicates is plotted for each genotype with error bars only going upward. Strains included are yRK114, yRK128, yRK104, and yRK92. (D) Plasmid end-joining assay results with three technical replicates for each of two biological replicates. Black circles correspond to the first biological replicate and pink triangles correspond to the second biological replicate. The number of linear DNA transformation colonies for each genotype relative to the average number of circular DNA transformation colonies for that genotype. An unpaired two-tailed student *t*-test was performed between the samples. Comparison of *MRX-tag* to *mrx-18A* had a p-value = 0.0.0680 and was not significant (n.s.). *MRX-tag* was not significantly different from wildtype (data not shown). Comparison of *mrx-18A* to *mre11Δ* had a p-values < 0.0001 (***). Strains included are segregants from yRK79, yRK80, yRK81, yRK83, and yRK5064. (E) Southern blot analysis of telomeres from strains with the indicated genotype. Two independent, haploid segregants were assayed for each genotype. The *mre11Δ* haploids were yRK1018 and yRK1019 and were passaged for approximately 200 population doublings. *WT*, *MRX-tag*, and *mrx-18A* haploids were segregants from yRK79, yRK80, yRK81, and yRK83. (F) Southern blot analysis of telomeres from strains with the indicated genotype. Haploid cells were passaged on solid media for approximately 140 population doublings. Passages 1, 3 and 6 are shown for simplicity. Two independent haploids segregants were assayed for each genotype. There were no growth defects observed over time. Segregants are from JHUy868, yRK79, yRK80, yRK81, and yRK83.

To examine the role of Tel1 and Mec1 phosphorylation of MRX on the DNA damage response, we tested the *mrx-18A* mutant for MMS sensitivity. None of the MRX S/T-Q mutants individually or in combination showed increased sensitivity to MMS, hydroxyurea, bleomycin, or 4-nitroquinoline (Figure S3A and data not shown). We noted *tel1Δ* alone does not have detectable MMS sensitivity (Figure S3A) but there is an observable increase in MMS sensitivity in *tel1Δ mec1Δ sml1Δ* compared to *mec1Δ sml1Δ* (Figure S3B). To take advantage of this increased sensitivity, we put *mrx-18A* and *MRX-tag* in the sensitized background of *mec1Δ sml1Δ* and assayed cells for an increased MMS sensitivity compared to *mec1Δ sml1Δ* alone. Using the spotting assay, it was difficult the determine whether the sensitivity of *mrx-18A mec1Δ sml1Δ* cells was increased compared to *MRX-tag mec1Δ sml1*Δ cells (Figure S3B). To quantify the subtle DNA damage defect of *mrx-18A mec1Δ sml1Δ* cells we used a more sensitive assay where colony forming units are counted as cells are treated with MMS over time. In this quantitative assay we observed no difference in MMS sensitivity between *mrx-18A mec1Δ sml1Δ* cells compared to *MRX-tag mec1Δ sml1*Δ (Figure 3C). We also noted in this assay that *MRX-tag mec1Δ sml1Δ* cells were slightly more sensitive than *mec1Δ sml1Δ*, suggesting the tags have a small effect on MRX complex function (Figure 3C). The epitope tags do not greatly disrupt function as this small effect was only observed in a *mec1Δ sml1*Δ sensitized background. No increased sensitivity was observed in response to 4-nitroquinoline, bleomycin, or hydroxyurea (Figure S4A-C). As a control, we also examined *mrx-18A tel1Δ,* and found no increased MMS-sensitivity (Figure S4D).

In addition to its role in homology-directed repair, MRX also plays a critical role in Non-Homologous End Joining (NHEJ) in yeast (MOORE AND HABER 1996). We tested the effect of *mrx-18A* on NHEJ using a plasmid re-ligation assay (BOULTON AND JACKSON 1996) and found no effect of *mrx-18A* compared to *MRX-tag* (Figure 3D). These data suggest that phosphorylation of MRX by on S/T-Q sites does not play a major role in NHEJ.

We next examined the telomere length phenotype of the *mrx-18A* mutant and found no effect compared to the *MRX-tag* cells. *MRX-tag* cells exhibited slight telomere shortening compared to untagged alleles, consistent with a slight defect seen in the DNA damage assay, however, *mre11Δ* was significantly shorter by comparison (Figure 3E). Individual mutant subunits, *mre11-4A, rad50-10A, and xrs2-4A,* also had no effect on telomere length (Figure S5). To determine whether a change in telomere length would be seen after further cell divisions, we streaked *mrx-18A* cells for approximately 120 population doublings and still saw no effect on telomere length (Figure 3F). These data suggest that phosphorylation of the MRX complex by Tel1 or Mec1 on S/T-Q sites is not critical for telomere length regulation.

### MRX is required downstream of Tel1 for the DNA damage response but not telomere length

The lack of requirement for MRX phosphorylation by Tel1 or Mec1 raised the question of whether MRX is required downstream of Tel1. Because double mutants of either *mre11Δ, rad50Δ, or xrs2Δ* with *tel1Δ* produces a short telomere phenotype similar to any individual mutant, it is not possible to establish epistasis. Therefore, we performed epistasis analysis with the *TEL1-hy909* hypermorphic allele and *mre11Δ, rad50Δ, or xrs2Δ*. As described earlier, *TEL1-hy909* elongates telomeres and provides a stark contrast to the short telomeres in *mre11Δ, rad50Δ, or xrs2Δ* cells. Many studies suggest Tel1 and MRX act in the same epistasis pathway and current models indicate that MRX recruits Tel1 to a DNA double-strand break, activates the kinase, Tel1 phosphorylates MRX, and then MRX processes DNA ends for repair (OH AND SYMINGTON 2018; PAULL 2018). Analogous pathways have been proposed for Tel1 and MRX in telomere length regulation, (TSUKAMOTO *et al*. 2001; VISCARDI *et al*. 2007; BONETTI *et al*. 2009) predicting that the MRX complex is downstream of Tel1 and that the MRX complex should be epistatic to Tel1 in both the DNA damage response and telomere length regulation.

To examine the epistasis of MRX and Tel1, we generated strains heterozygous for *TEL1/TEL1-hy909 MEC1/mec1Δ* and *SML1/sml1Δ* together with individual heterozygous deletions of *mre11Δ, rad50Δ, or xrs2Δ* and initially examined the DNA damage response. We observed the *TEL1-hy909 mre11Δ* double mutants were as sensitive to MMS as *mre11Δ* alone (Figure 4A), consistent with previous work (BALDO *et al*. 2008). The other two double mutants, *TEL1-hy909 rad50Δ* and *TEL1-hy909 xrs2Δ*, were also both MMS sensitive (not shown). These results support a role for MRX downstream of Tel1 in the DNA damage response, as previously reported (MOORE AND HABER 1996; USUI *et al*. 2001). We found that, surprisingly, *TEL1-hy909 mec1Δ mre11Δ* spores were inviable, indicating the rescue of *mec1Δ* lethality by *TEL1-hy909* is MRX-dependent (Figure S6). This was also true for *TEL1-hy909 mec1Δ rad50Δ* and *TEL1-hy909 mec1Δ xrs2Δ* (not shown).

**Figure 4.**
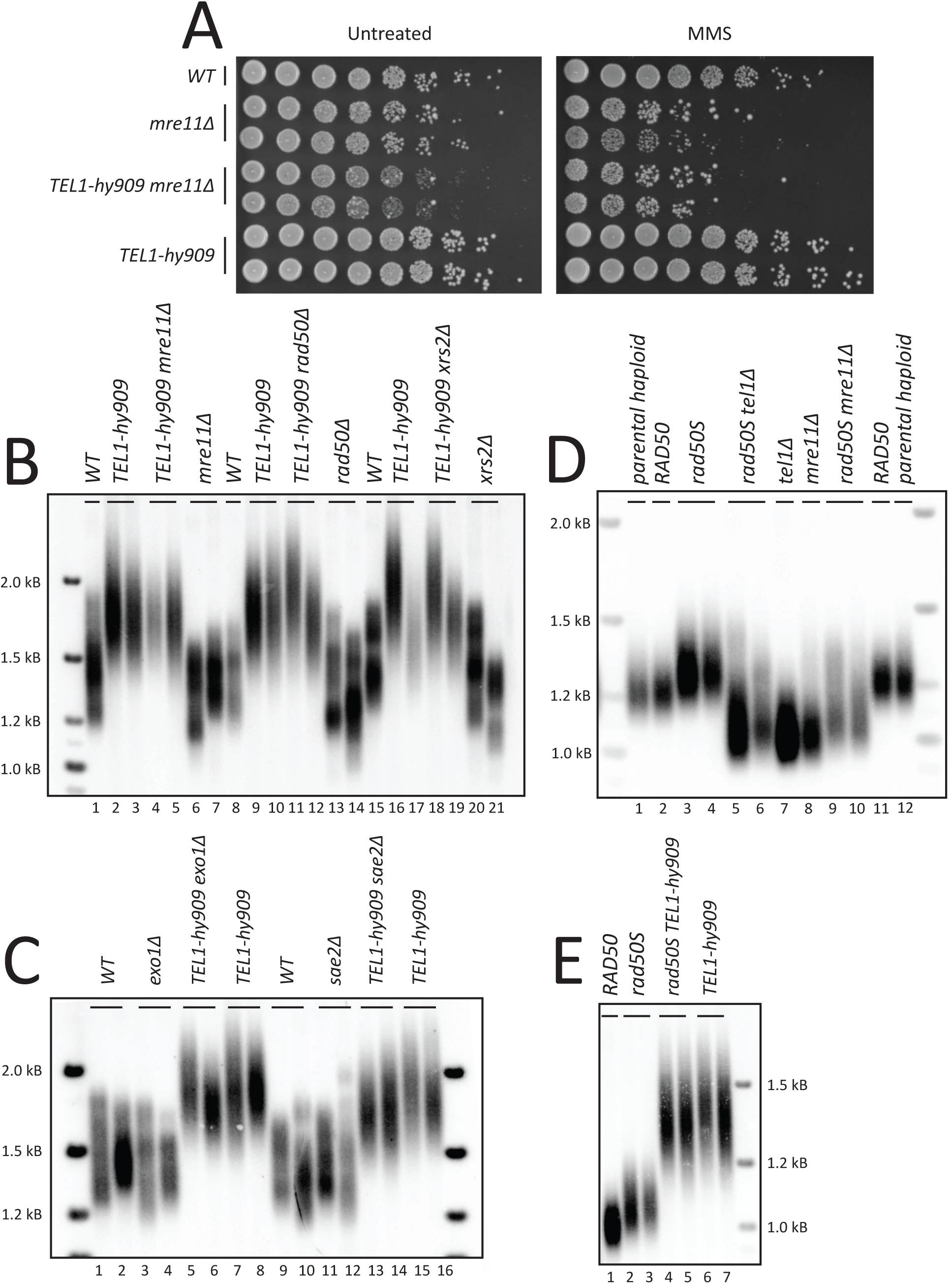
Tel1-hy909 requires the MRX complex for the DNA damage response but not for telomere elongation. (A) Yeast dilution series of untreated cells or cells treated with 0.02% MMS for one hour. The genotype is indicated to the left of the panels. To account for growth differences between the genotypes different amounts of cells were collected for the initial dilution. 0.5 OD of cells were collected for *WT* and *TEL1-hy909*, 1.5 OD of cells were collected for *mre11Δ,* and 8.0 OD of cells were collected for *TEL1-hy909 mre11Δ*. Strains used in this assay were yRK114, yRK126, yRK128, yRK104, yRK141, yRK92, yRK93, and yRK122. (B) Southern blot analysis of telomeres from strains with the indicated genotype. Two independent, haploid segregants were assayed for each genotype. Segregants are from JHUy816, yRK79, yRK80, yRK81, and yRK83. Cells underwent minimal propagation before genomic DNA was prepared. (C) Southern blot analysis of telomeres from strains with the indicated genotype. Two independent haploids segregants were assayed for each genotype. Cells underwent minimal propagation before genomic DNA was prepared. Segregants are from yRK5089, yRK5090, yRK5093, and yRK5094. (D)-(E) CRISPR/Cas9 was used to knock-in the *rad50S* allele into a wild-type haploid strain (yRK114). A transformant that was not edited at the *RAD50* locus but was transformed with the Cas9 plasmid was used as a control and is referred to as *RAD50* (Figure 5D, lane 2). Both *RAD50* and *rad50S* transformants were passaged on solid media for approximately 120 population doublings (see 5D lanes 2-4, yRK2112-5, yRK2113-5, and yRK2116-5). *rad50S* or *RAD50* cells were transformed to introduce *tel1Δ, mre11Δ* (Figure 5D lanes 5-10), or *TEL1-hy909* (Figure 5E lanes 4-7). Cells were passaged on solid media for approximately 120 population doublings. The strains used were yRK2118-5, yRK2120-5, yRK2121-5, yRK2122-5, yRK2123-5, yRK2124-5, yRK2125-5, yRK2126-5, yRK2127-5, and yRK2128-5.

We next examined telomere length in *TEL1-hy909 mre11Δ, TEL1-hy909 rad50Δ,* and *TEL1-hy909 xrs2Δ* double mutants and found that, surprisingly, in all three cases the double mutants had long telomeres, similar to *TEL1-hy909* alone (Figure 4B). This indicates that, unlike the DNA damage response, MRX is not epistatic to Tel1 for telomere length. Because this result differs from previous findings, we repeated the experiment in the strain background (W303) used in the previous study (BALDO *et al*. 2008). In this analysis, all three independently derived double mutants *TEL1-hy909 mre11Δ, TEL1-hy909 rad50Δ,* and *TEL1-hy909 xrs2Δ* showed long telomeres consistent with our initial findings (Figure S7A). We also passaged *TEL1-hy909 xrs2Δ* for 120 population doublings to see if shortening might occur with further divisions. Instead, we found telomeres elongated further with passaging (Figure S7B). These data suggest that MRX is not required downstream of Tel1-hy909 to carry out telomere elongation.

To further investigate the requirement of nuclease activity at the telomere, we tested the epistatic relationship between *TEL1-hy909* and deletion of Sae2(CtIP), which stimulates Mre11 (LENGSFELD *et al*. 2007), or Exo1, which has been suggested to play a role in telomere processing (MOREAU *et al*. 2001). We generated diploid strains that were heterozygous for *TEL1/TEL1-hy909 EXO1/exo1Δ* and diploid strains that were heterozygous for *TEL1/TEL1-hy909 SAE2/sae2Δ*. *TEL1-hy909 exo1Δ* cells and *TEL1-hy909 sae2Δ* cells had long telomeres, similar to *TEL1-hy909* alone (Figure 4C, compare lanes 5-6 to lanes 7-8 and lanes 13-14 to lanes 15-16). These data support the hypothesis that telomere resection by MRX/Sae2 or Exo1 is not required for telomere length maintenance.

### Rad50S activates Tel1 for telomere length maintenance

The ability of Tel1-hy909 to generate long telomeres in the absences of MRX suggests that this hypermorph is constitutively active as it does not require activation by MRX. As an independent approach to determine whether MRX is only required upstream of Tel1 in telomere length regulation, we performed the a similar epistasis experiment using *tel1Δ* and a previously identified MRX complex mutant, *rad50S* (ALANI *et al*. 1990). Rad50S produces a long telomere phenotype which has been attributed to increased Tel1 activation (KIRONMAI AND MUNIYAPPA 1997). *rad50S* mutants are reported to have a sporulation defect (USUI *et al*. 2001), therefore we performed these experiments in haploid cells. We used CRISPR/Cas9 to knock-in the *rad50S* allele at the endogenous *RAD50* locus (Anand et al. 2017). Elongated telomeres were observed after cells were passaged for approximately 120 population doublings (Figure 4D, lanes 3-4). In these haploids, we subsequently introduced a *tel1Δ* or *TEL1-hy909* allele at the *TEL1* locus. As a control, parallel strains were generated where a *mre11Δ* allele was introduced at the *MRE11* locus. Without Mre11, Rad50S should not be able function in the MRX complex and the hypermorph activity will not be observed. *rad50S tel1Δ* double mutants showed short telomeres, similar in length to *tel1Δ* (Figure 4D) indicating *rad50S* does not affect telomere elongation by acting downstream of Tel1. Telomere shortening was also observed in *rad50S mre11Δ* cells, as expected (Figure 4D). These experiments suggest that Tel1 is required for the telomere elongation seen in *rad50S* cells. Further, *rad50S TEL1-hy909* had very long telomeres similar to *TEL1-hy909* (Figure 4E). Our data indicate that MRX activates Tel1 but does not contribute to processing of telomeres to allow telomere length regulation after Tel1 activation. In contrast, the MRX complex is critical downstream of Tel1 for the DNA damage response (Figure 5).

**Figure 5.**
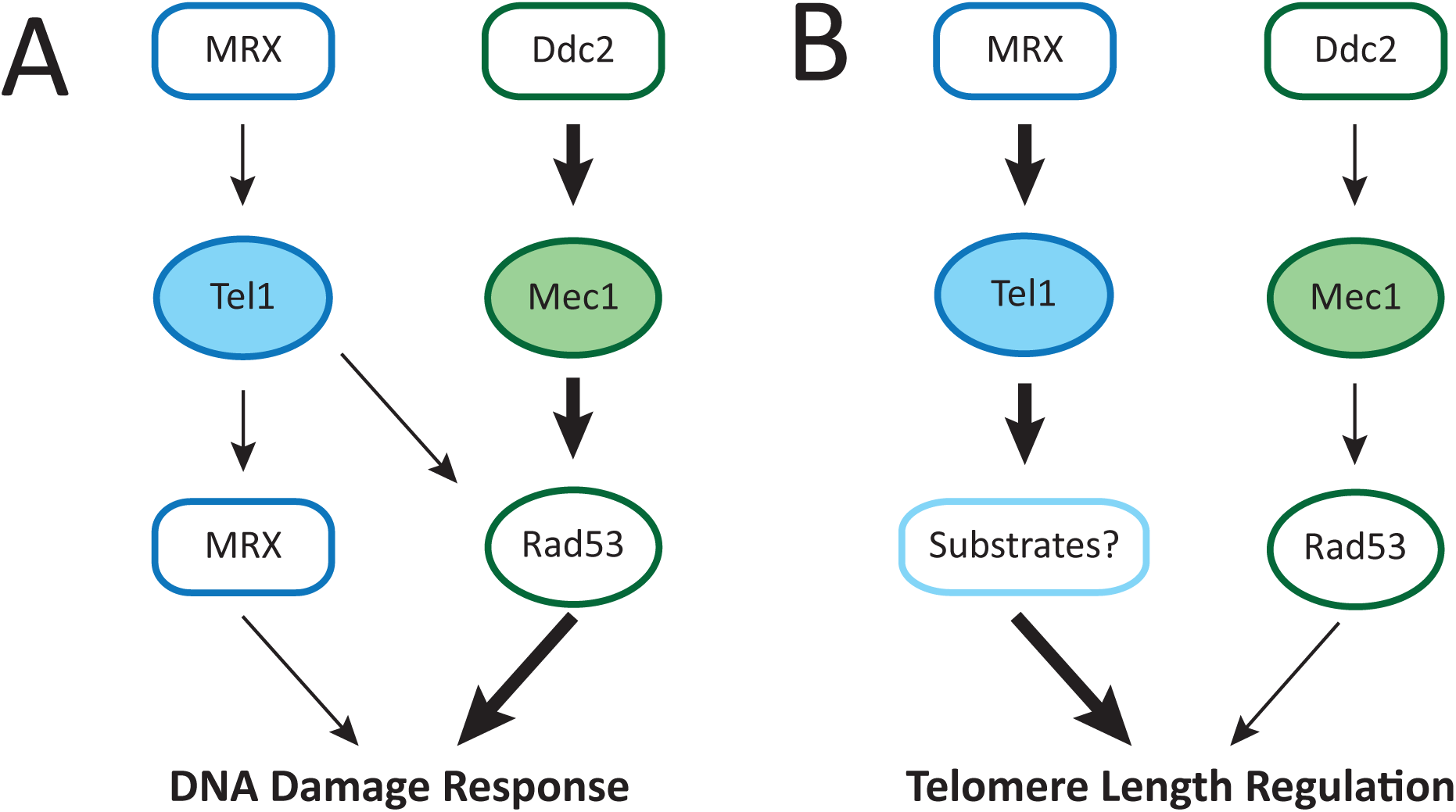
Tel1 regulates telomere length in a pathway distinct from the DNA damage response. Diagram demonstrating the distinctions between Tel1 pathways in the DNA damage response and telomere length regulation. (A) The DNA damage response is most strongly regulated by Mec1 and Rad53 as indicated with the bold arrows, although Tel1 signaling through Rad53 and MRX plays a role. The MRX complex is both upstream and downstream of Tel1 in the DNA damage response. (B) For telomere length regulation, Tel1 does not require MRX after activation and Rad53 does not plan a role in the Tel1 telomere length regulation pathway. The Tel1/MRX pathway plays the major role in telomere length compared to a minor role of Mec1/Rad53 pathway.

## Discussion

### Telomere elongation and DNA damage response are regulated through different mechanisms

We found that, in contrast to published models, the MRX complex is not required after Tel1 activation for telomere elongation. Our data suggest a new model for the regulation of telomere length in which Tel1 activation by MRX is sufficient for telomere length regulation but not for the DNA damage response (Figure 5). Previous work has shown that Mec1 and Rad53 play a major role in DNA damage response, while Tel1 plays a minor role acting through Rad53 and the MRX complex. In the DNA damage response, the MRX complex is thought to act both upstream and downstream of Tel1 (USUI *et al*. 2001; PAULL 2015). MRX binds to double-strand breaks and interacts with Tel1, activating its kinase activity. Previously, a parallel model for telomere length regulation suggested that MRX recruits and activates Tel1 at the telomere, then MRX processes telomere ends to promote telomerase elongation (LARRIVEE *et al*. 2004; BONETTI *et al*. 2009). In this model, Mec1 is considered secondary to Tel1 for telomere length regulation and its function was presumed to be redundant. In contrast, we show that Mec1 and Rad53 act in a separate, non-overlapping pathway from Tel1 for telomere length maintenance. Together these data demonstrate that the Tel1 and Mec1 pathways differ significantly for the DNA damage response and telomere length regulation.

### Dysregulation of dNTP pools can mask telomere length phenotypes

The role of Mec1 in telomere length regulation has remained poorly understood, in part because of discrepancies in reported telomere length phenotypes. *mec1Δ sml1Δ* telomeres appear similar to wildtype while *mec1Δ crt1Δ* telomeres are shorter than wildtype (Figure 1B). Both *sml1Δ* and *crt1Δ* suppress the lethality of *mec1Δ* through upregulation of different pathways that regulate nucleotide pools (HUANG *et al*. 1998; ZHAO *et al*. 1998). Several studies suggest that the increased telomere length in *mec1Δ sml1Δ* compared to *mec1-1* and *mec1-21* alleles is due to increased telomerase processivity with increased dGTP levels (GUPTA *et al*. 2013). Recent work has suggested that while both mutants increase nucleotide pools, *sml1Δ* and *crt1Δ* have different effects on the specific ratio of dGTP to other dNTPs (MAICHER *et al*. 2017). Because dGTP is limiting for telomerase processivity *in vitro* (GREIDER AND BLACKBURN 1987; HAMMOND AND CECH 1997), it was proposed that an increased dGTP/dNTP ratio would elongate telomeres. (MAICHER *et al*. 2017). However, increased dNTP levels are not sufficient to lengthen telomeres, as *sml1Δ* and *crt1Δ* mutants do not show increased telomere length on their own. Also, *crt1Δ* does not lengthen telomeres in either a *tel1Δ* or *mec1Δ* background (Figure 1B). Therefore, while changes in dNTP pools in *mec1Δ sml1Δ* cells may mask telomere length phenotypes (LONGHESE *et al*. 2000) the data are not consistent with altered telomerase processivity as the mechanism.

### Rad53 phosphorylation by Mec1 contributes to telomere length regulation

Previous work has shown that Tel1 or Mec1 phosphorylation of Rad53 is critical for the DNA damage response. Our data demonstrate an additional role for Rad53 phosphorylation in telomere length regulation. This phosphorylation is likely primarily performed by Mec1, as our data indicate that Rad53 is in the Mec1 telomere length pathway and it has previously been shown that Mec1 phosphorylation of Rad53 is predominant in the DNA damage response (USUI *et al*. 2001). We cannot exclude the possibility that Tel1 phosphorylation of Rad53 contributes in a small way to telomere length regulation. However, the *TEL1-hy909* hypermorphic allele showed telomere elongation in the absence of Rad53 (Figure 2D), suggesting that Tel1 does not require Rad53 for telomere length regulation. Our model suggests there are as yet unknown substrates that mediate the Tel1 effect on telomere length (Figure 5).

### Rad53 is a critical mediator of Mec1 in telomere length regulation

The *TEL1-hy909* hypermorphic allele can rescue the lethality of *mec1Δ,* as shown previously (BALDO *et al*. 2008). However, we found that *TEL1-hy909* did not rescue *rad53Δ* lethality. Rad53 is a substrate of both Tel1 and Mec1 (SANCHEZ *et al*. 1996; SMOLKA *et al*. 2007). Both *mec1Δ* and *rad53Δ* are thought to be lethal due to an inability to upregulate ribonucleotide reductases for DNA repair. Tel1-hy909 has increased catalytic activity *in vitro* and is able to phosphorylate Rad53 more efficiently than Tel1 (BALDO *et al*. 2008). Tel1-hy909 likely rescues *mec1Δ* lethality because of its increased ability to activate Rad53. Our finding that *TEL1-hy909* cannot rescue *rad53Δ* places Rad53 as the critical mediator of Mec1: Mec1 loss can be compensated for by Tel1-hy909 but this hypermorph cannot compensate for loss of Rad53.

The *MRX* complex is epistatic to *TEL1-hy909* in the DNA damage response, as *TEL1-hy909 mre11Δ* was just as sensitive as *mre11Δ* to MMS challenge. This was also true for the other MRX complex components. We unexpectedly found that *TEL1-hy909 mec1Δ mre11Δ* is lethal while *TEL1-hy909 mec1Δ* is viable (Figure S6). It is unclear why the *TEL1-hy909* rescue of *mec1Δ* viability is *MRX*-dependent. Previous models would suggest this is because the MRX complex is required for Tel1 activation. However, our data indicate that the *TEL1-hy909* allele is constitutively active.

### MRX complex phosphorylation by Tel1/Mec1 on S/T-Q sites is not required for DNA damage response, NHEJ, or telomere length regulation

Multiple studies have reported that Tel1/Mec1-dependent phosphorylation of the MRX complex occurs in response to DNA damage. Thus, were surprised to find that the mrx-18A mutant did not have a DNA damage phenotype or in NHEJ. The absence of an effect on telomere length was also surprising, and suggests MRX is not the substrate that mediates the Tel1 pathway of telomere length regulation.

### Telomere elongation can occur in the absence of MRX complex

The fact that Tel1-hy909 telomere elongation can occur in the absence of the MRX complex indicates that telomere elongation is possible without telomere end processing by MRX. This epistasis indicates that the Tel1-hy909 hypermorph has bypassed the need to interact with MRX for its activation, and that the point mutations in the *TEL1-hy909* allele promote constitutive catalytic activity. This finding, combined with the fact that the mrx-18A mutant has no telomere length defect, suggest that Tel1 does not require MRX for telomere length regulation after it is activated. Further, the only known mutants in MRX that decrease telomere length are those that decrease the MRX complex interaction with Tel1. (i.e. Xrs2 C-terminal truncation) (NAKADA *et al*. 2003a; MA AND GREIDER 2009). Similarly, mutants that increase telomere length are thought to hyperactivate Tel1 (i.e. *rad50S* and *TEL1-hy* alleles) (KIRONMAI AND MUNIYAPPA 1997; BALDO *et al*. 2008). MRX mutants that alter the catalytic functions of the complex are required for the DNA damage response but not for telomere length regulation. For example, alleles of Mre11 that lack nuclease function do not show a telomere length phenotype (MOREAU *et al*. 1999; TSUKAMOTO *et al*. 2001) but do inhibit the DNA damage response (BUIS *et al*. 2008). Similarly, deletions of the Mre11 nuclease co-factor, Sae2 (CtIP) alone, or in combination with Exo1 deletion, show no telomere length defects (BONETTI *et al*. 2009). This is consistent with our observation that, like *mre11Δ, TEL1-hy909* elongates telomeres in *sae2Δ* and *exo1Δ* cells (Figure 4C). We conclude that cells with deletions of MRX complex components have short telomeres because of the reduction in Tel1 activation, not because the cell lacks the resection functions.

Our finding that MRX is not required downstream of Tel1 for telomere elongation has important implications for telomere elongation models. Most models suggest that after replication of the telomere the leading strand telomere is processes by a nuclease before telomerase can elongate it. The presumption that leading strand replication leaves a blunt end that requires processing is an assumption that has not been directly tested (LINGNER AND CECH 1998; PFEIFFER AND LINGNER 2013). In contrast to those models, our data suggest telomerase can efficiently elongate telomeres without end processing. Tel1(ATM) and the MRX(N) complex are thought to function by similar mechanisms in *S. cerevisiae* and mammalian cells (OH AND SYMINGTON 2018; PAULL 2018). Therefore, the data presented here suggest we should rethink the requirements for telomere resection preceding telomere elongation broadly across all organisms.

## Author Contributions

Study conception: CWG and RK, Methodology: RK and CWG, Investigation and Acquisition of data: RK and CC, Supervision: CWG, Formal analysis and interpretation of data: RK and CWG, Validation: RK and CC, Visualization: RK, Data curation: RK, Funding acquisition: CWG, Drafting of manuscript: RK and CWG, Critical revision: RK, CC, and CWG

## Acknowledgements

We thank Rini Mayangsuri for technical assistance with this work. We thank Brendan Cormack, Geraldine Seydoux, the Greider Lab, and the Armanios Lab for critical reading of the manuscript and helpful discussions. This work was supported by Bloomberg Distinguished Professorship (to CWG) and the Turock Fellowship (to RK).

